# Embryo geometry drives formation of robust signaling gradients through receptor localization

**DOI:** 10.1101/491290

**Authors:** Zhechun Zhang, Steven Zwick, Ethan Loew, Joshua S. Grimley, Sharad Ramanathan

## Abstract

Morphogen signals are essential for cell fate specification during embryogenesis. Some receptors that sense these morphogen signals are known to localize to only the apical or basal membrane of polarized cell lines *in vitro*. How such localization affects morphogen sensing and patterning in the developing embryo remains unknown. Here, we show in the early mouse embryo that the formation of a robust BMP signaling gradient depends on restricted, basolateral localization of the BMP receptors. Mis-localizing these receptors to apical membrane leads to ectopic BMP signaling *in vivo* in the mouse embryo. To reach the basolaterally localized receptors in epiblast, BMP4 ligand, secreted by the extra-embryonic ectoderm, has to diffuses through the narrow interstitial space between the epiblast and the underlying endoderm. This restricted, basolateral diffusion creates a signaling gradient. The embryo geometry further buffers the gradient from fluctuations in the levels of BMP4. Our results demonstrate the importance of receptor localization and embryo geometry in shaping morphogen signaling during embryogenesis.

## Introduction

Morphogens are long-range signaling molecules, that move in extracellular space to induce concentration-dependent cellular responses in their target tissues ^1,2^. Genetic perturbation of morphogens and their cognate receptors often leads to missing cell types and embryonic structures ^3–6^. Many mechanisms have been proposed to explain how morphogens induce signaling gradients in target tissues and therefore direct the spatial organization of cell fates ^1,2,7–15^. Surprisingly, morphogen receptors can localize to either apical or basolateral membrane of the epithelial tissues. Such localization can dramatically affect how the target tissue senses morphogens. How receptor localization modulates morphogen signaling in developing embryos is not known.

The early mouse embryo (E6.0-E6.5) adopts an egg-cylinder geometry (Fig. 1a) ^5,6,16^. It contains a lumen (the pre-amniotic cavity) encased by two epithelial tissues: the epiblast and extra-embryonic ectoderm (ExE). The ExE secretes the morphogen BMP4, which is sensed by receptors in epiblast ^5,6^. The resulting BMP signaling is required for the differentiation of the epiblast into mesoderm ^3,4^

**Fig. 1.**
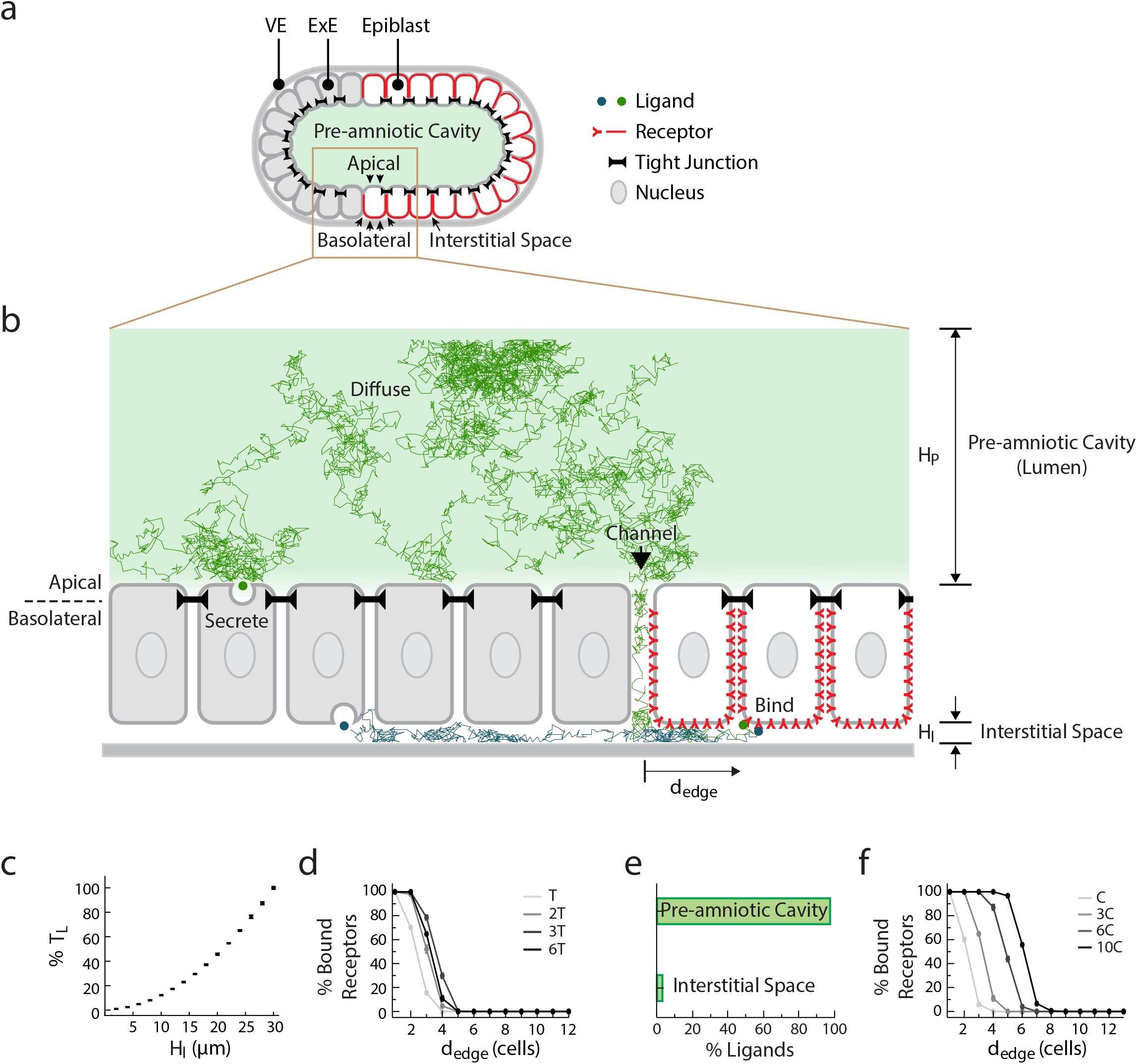
Receptor localization facilitate the formation of a robust signaling gradient in early mouse embryo. **a**, Illustration of pre-gastrulation mouse embryo, with the epiblast (white) and extraembryonic ectoderm (ExE, light grey) together enclosing the pre-amniotic cavity. Apical membranes of epiblast cells face the pre-amniotic cavity whereas basolateral membranes face the interstitial space. **b**, Illustration of a simulation with basolateral receptors. ExE cells (light gray) secrete BMP4 ligands from their apical or basolateral membranes, while epiblast cells have BMP receptors (red) on their basolateral membranes. Ligands cannot diffuse past tight junctions between cells (black). Simulated ligand trajectories show that ligands diffuse from epiblast edge (black arrow) through interstitial space to approach and bind basolateral receptors. **c**, T_L_ (the time between BMP4 ligands entering interstitial space and being captured by receptors), increases with the height of interstitial space, H_I_. **d**, Plot of percentage of ligand-bound receptors as a function of their distance from epiblast edge, d_edge_, over time in simulations with apically secreted ligands (T=5 min). **e**, Percentage of unbound ligands in pre-amniotic cavity vs. interstitial space at steady state (6T) in simulations with apically secreted ligands. **f**, Plot of percentage of ligand-bound receptors as a function of d_edge_ at steady state (6T) shows signaling gradients at different BMP4 concentrations in simulations with apically secreted ligands (C=0.08 ng/mL).

Both the epiblast and ExE have stereotyped epithelial tissue geometries ^17^, with their apical membranes surrounding the lumen and their basolateral membranes facing a narrow interstitial space (between these tissues and the underlying visceral endoderm (VE)). This lumen and interstitial space are separated by impermeable tight junctions present throughout the epithelia except at the border between the ExE and epiblast (Fig. 1a). Indeed, when small-molecule dye fluorescein was injected into the pre-amniotic cavity of an E6.5 mouse embryo, it did not penetrate the epiblast or ExE but diffused through a channel at the edge of the epiblast (Supplementary Fig. 1). Thus, the extracellular space in the embryo through which BMP4 ligands diffuse is compartmentalized into a lumen and an interstitial space.

Here, by combining mathematical modeling, quantitative imaging, embryological perturbation, and microfluidics, we demonstrate that restricted receptor localization in conjunction with the embryo geometry, constrains the diffusion of and therefore response to BMP4 ligands. We show that the BMP4 signaling gradient arises from the edge of epiblast even under conditions of uniform BMP4 stimulation. Further, the interplay between restricted receptor localization and the embryo geometry buffers BMP4 ligands in pre-amniotic cavity through an entropic effect. This entropic buffering renders the BMP4 signaling gradient robust to fluctuations in BMP4 level. Consistently, mis-localizing BMP receptors in the mouse embryo leads to ectopic BMP4 signaling. Thus, receptor localization and embryo geometry together play an essential role in regulating morphogen signaling during early development.

## Results

### Receptor localization facilitates the formation of a robust signaling gradient in early mouse embryo

To understand how receptor localization impacts BMP signaling between the ExE and epiblast, we simulated the movement of individual BMP4 ligands in early mouse embryo (E6.0-E6.5) from secretion to receptor binding, using Langevin dynamics ^18^. Given the evidence of polarized ligand secretion by epithelial cells *in vitro* ^19,20^, we modeled different instances in which BMP4 ligands were secreted apically (into the pre-amniotic cavity) or basolaterally (into the interstitial space) by the ExE (Fig. 1b). After secretion, ligands diffused through extracellular space in the embryo. Due to tight junctions in the simulation, ligands could move between the pre-amniotic cavity and the interstitial space only by diffusing through the channel between the ExE and epiblast. Some signaling receptors are known to localize to only the apical or basolateral membranes of epithelial cells ^14,20–23^; such localization could determine the compartment from which ligands are sensed by receptors in the epiblast. Therefore, we also performed simulations with BMP receptors localized exclusively on the apical membrane (facing the pre-amniotic cavity) or the basolateral membranes (facing the interstitial space) of the epiblast cells. Finally, our model assumed that once BMP4 ligands bound their receptors, signaling activity was induced and the ligands were cleared.

Our simulations revealed that if the BMP receptors are basolaterally localized in the epiblast, the compartmentalized geometry of the embryo naturally results in the formation of a robust BMP signaling gradient. This occurs despite the absence of other regulatory mechanisms such as signaling inhibitors ^2,9–11,24^ The basolateral localization of BMP receptors requires that ligands diffuse through the interstitial space between the epiblast and VE to access them (Fig. 1b). The height of this interstitial space, HI, regulates the time, TL, and hence the distance a ligand can diffuse before being captured by a receptor (Fig. 1c). As a consequence, BMP4 ligands are more likely to bind receptors that are closer to the epithelial edge, giving rise to a BMP signaling gradient from the edge of the epiblast inward (Fig. 1d). The signaling gradient forms regardless of whether BMP ligands are secreted from the apical or basolateral membrane of the ExE and arises even if ligands are imposed to be uniformly distributed in the pre-amniotic cavity (Supplementary Fig. 2a).

The basolateral localization of BMP receptors, in conjunction with, the asymmetric compartmentalization of the embryo also makes this BMP signaling gradient robust to fluctuations in the BMP4 source strength. Due to the large volume difference between the pre-amniotic cavity and the interstitial space, and the channel (between ExE and epiblast) that connects these two compartments, the majority of BMP4 ligands accumulate in the cavity on the apical side of the epiblast. This is an entropic effect: the entropy of BMP4 ligands is increased as more ligands reach the pre-amniotic cavity. In other words, accumulation of BMP4 ligands in the cavity, is driven by the same physical forces, that allows ink to diffuse through water and ultimately reach uniform distribution independent of where ink is dropped initially. Consistently, BMP4 ligands accumulate in the pre-amniotic cavity, regardless of whether the ligands are secreted apically or basolaterally from the ExE in the simulation (Fig. 1e and Supplementary Fig. 2b).

This accumulation results in an entropic buffering effect: the pre-amniotic cavity serves as a ligand reservoir that buffers the signaling gradient against fluctuations in the BMP source strength. Indeed, if the total ligand concentration is increased by tenfold in a simulation with basolateral receptors, the signaling gradient shifts inward by only a few cell widths (Fig. 1f and Supplementary Fig. 2c). In striking contrast, if BMP receptors are apically localized in the epiblast or if tight junctions are absent, this ten-fold increase is sufficient to saturate all receptors in the simulation and destroy the signaling gradient (Supplementary Fig. 2c). Thus, the robustness of the BMP signaling gradient relies upon both the receptor localization and embryonic geometry in our simulation.

Assuming that the BMP receptors are basolaterally localized, our model provides three experimentally testable predictions. First, a BMP signaling gradient will form inward from the epiblast edge even if ligands are present at high concentration throughout the lumen (Supplementary Fig. 2a). Second, this signaling gradient will be robust to fluctuations in BMP concentration (Fig. 1f). Third, the mis-localization of BMP receptors to the apical membrane should lead to ectopic BMP signaling in the epiblast, since apically localized receptors will be able to detect BMP4 ligands that are buffered in the lumen (Fig. 1e).

### BMP receptors localize at the basolateral membrane of hESCs *in vitro* and mouse epiblast *in vivo*

We first asked whether BMP receptors are indeed basolaterally localized in mammalian cells. We measured the localization of these receptors through surface immunostaining ^21,23^ as well as by imaging GFP- and epitope-tagged receptors (see METHODS). The BMP co-receptors BMPR1A (Fig. 2b,c and Supplementary Fig. 4e-h) and BMPR2 (Supplementary Fig. 4i,j) are basolaterally localized in human embryonic stem cells (hESCs ^14^). We moreover found that the majority of TGF-β family receptors (including BMP receptors) in sequenced vertebrates contain a conserved LTA amino acid motif near their C-terminus (Fig. 2a and Supplementary Fig. 3). This motif has been shown to be necessary and sufficient for the basolateral localization of TGFBR2 in epithelial Madin-Darby canine kidney (MDCK) cells, and the mutation of this motif to an LTG sequence leads to the receptor’s mis-localization to the apical membrane ^21^. Consistently, we found that TGFBR2 and its co-receptor TGFBR1 are localized at the basolateral membrane of epithelial human hESCs (Supplementary Fig. 4a-d). Furthermore, the ACTIVIN/NODAL receptors ACVR1B and ACVR2B have also been found to be basolaterally localized in studies using human gastruloids ^14^, consistent with the fact that these receptors have LTA motifs (Supplementary Fig. 3). Thus, an evolutionarily conserved LTA motif is present in all of these receptors that are exclusively localized along the basolateral membrane in hESCs.

**Fig. 2.**
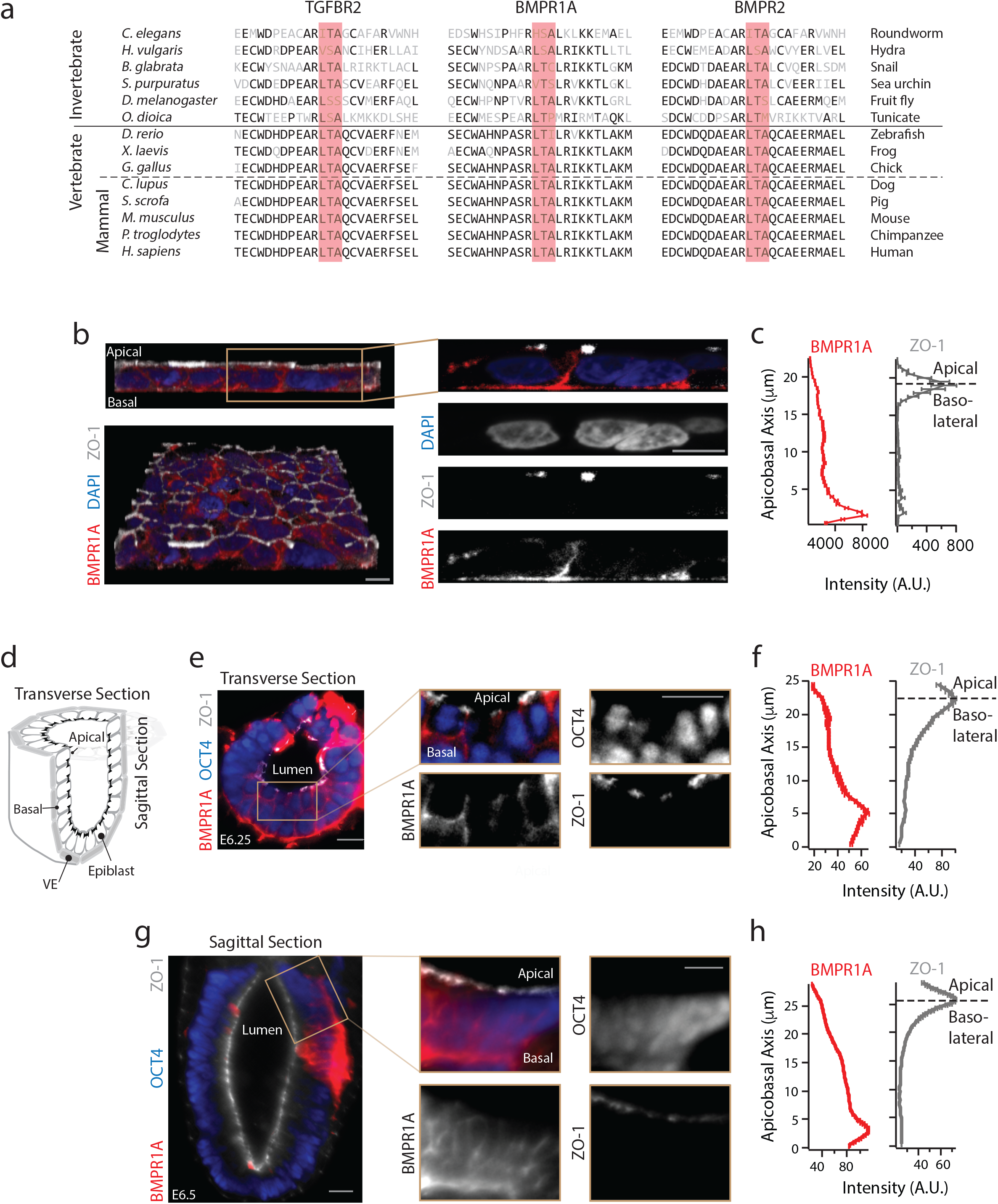
BMP receptors localize at basolateral membrane of hESCs *in vitro* and mouse epiblast *in vivo*. **a**, Protein sequence alignment of TGF-β and BMP receptors show the presence of LTA motif in receptors from multiple invertebrate and vertebrate species. **b**, *Left column*: 3D confocal images of hESC colony stained for BMPR1A (red), tight junction marker ZO-1 (white), and DNA (blue), in lateral (top) and tilted view (bottom). *Right column*: Zoomed-in lateral images. Scale bar 10 μm. **c**, Plots of BMPR1A (left) and ZO-1 (right) staining intensity along apicobasal axis show BMPR1A localized beneath tight junctions (n=38 cells). Error bars denote SEM. **d**, Illustration of pre-gastrulation mouse embryo, showing transverse and sagittal sections. **e**, Transverse section of an E6.25 mouse embryo stained for epiblast marker OCT4, BMPR1A, and ZO-1. **f**, Plots of BMPR1A (left) and ZO-1 (right) staining intensity along apicobasal axis for transverse section from (e) shows BMPR1A localized beneath tight junctions. **g,h**, Same as in (**e,f**) but for a sagittal section of an E6.5 mouse embryo.

**Fig. 3.**
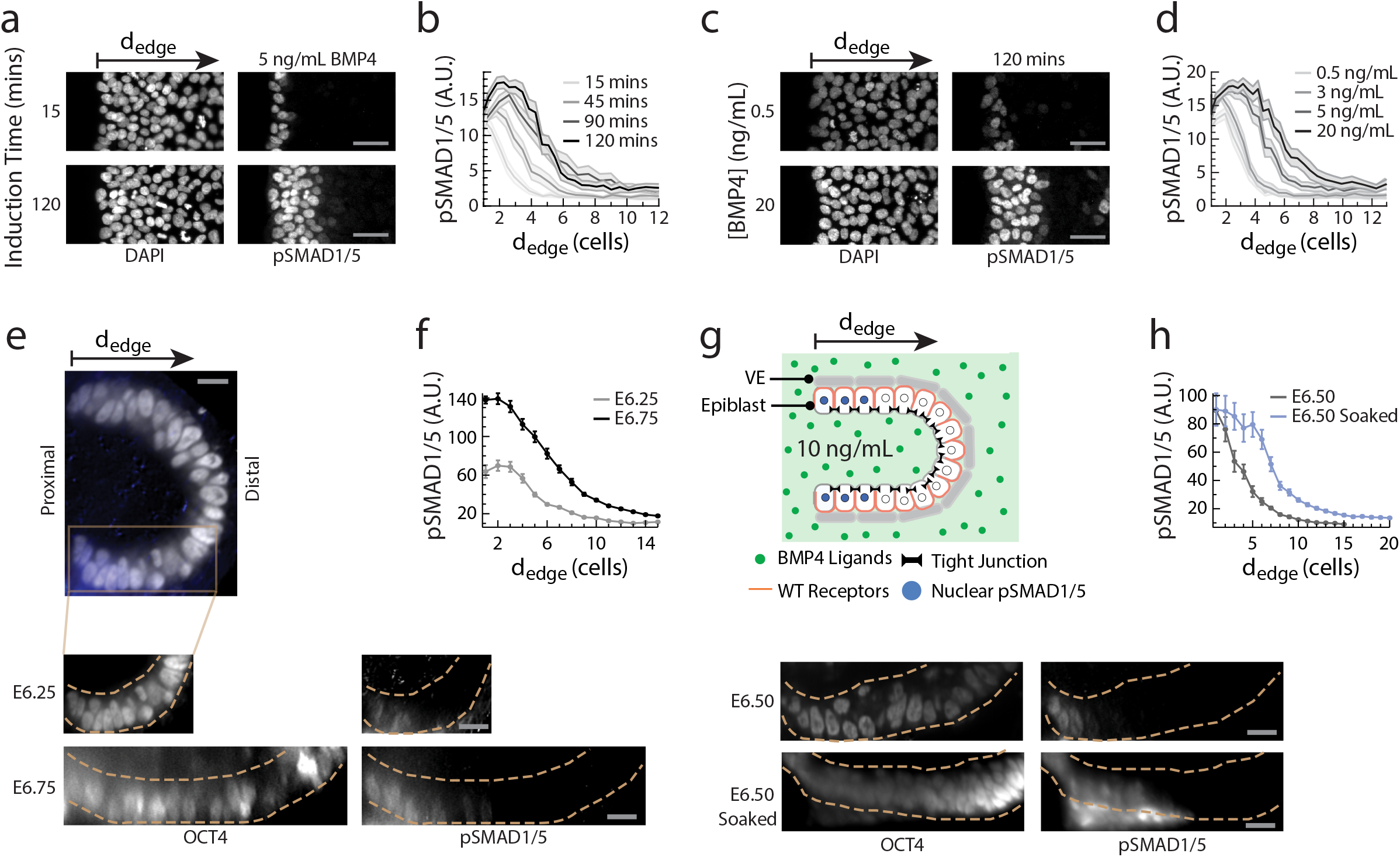
A robust BMP signaling gradient forms from the epiblast edge. **a**, hESC colony stained for DNA and pSMAD1/5 after 15 or 120 min BMP4 induction. **b**, pSMAD1/5 level of single hESCs as a function of their distance from the nearest colony edge, or d_edge_, after 15-120 min of BMP4 induction. **c**, hESC colony stained for DNA and pSMAD1/5 after 120 min exposure to 0.5-20 ng/mL BMP4. **d**, pSMAD1/5 level of single hESCs as a function of d_edge_ after 120 min exposure to 0.5-20 ng/mL BMP4. **e**, E6.25-E6.75 mouse embryos stained for OCT4 and pSMAD1/5. **f**, pSMAD1/5 level of mouse epiblast cells as a function of their distance from the posterior proximal edge of the epiblast, or d_edge_, for E6.25 and E6.75 embryos. Dotted yellow lines indicate epiblast boundary. **g**, *Top:* Illustration of BMP4 exposure experiment. ExE is surgically removed, and remaining epiblast-VE cup is soaked in media containing 10 ng/mL BMP4 for 30 min. *Bottom*: Intact E6.5 mouse embryo and BMP4-soaked E6.5 mouse embryo, both stained for OCT4 and pSMAD1/5. Dotted yellow lines indicate epiblast boundary. **h**, pSMAD1/5 intensity of epiblast cells as a function of d_edge_ for intact and BMP4-exposed E6.5 embryos. In all mouse data, error bars denote SEM and scale bar 20 μm.

**Fig. 4.**
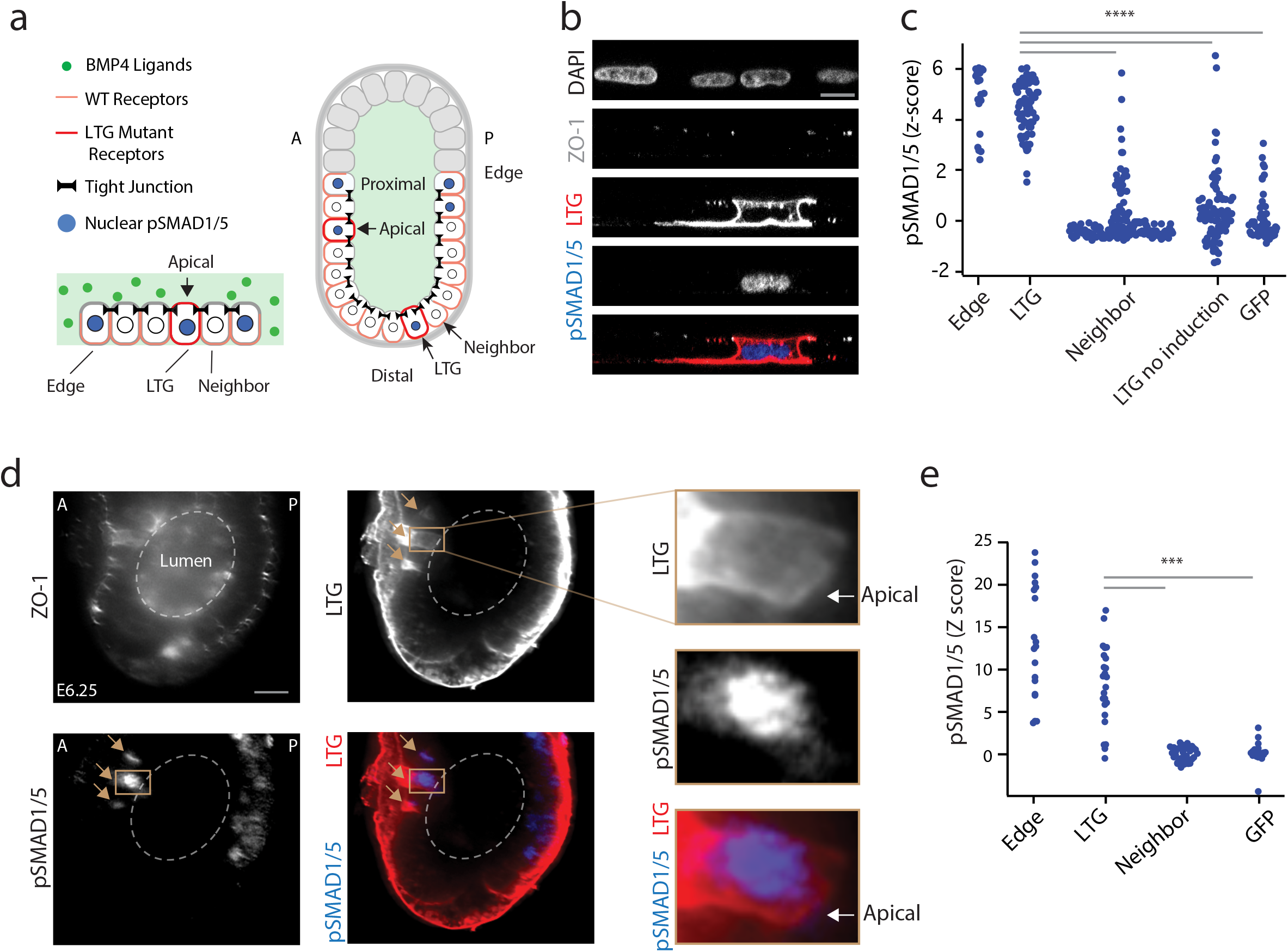
Mis-localization of receptors leads to ectopic BMP4 signaling. **a**, *Left:* Illustration of a hESC colony containing a single cell with mis-localized LTG mutant BMP receptors, exposed to BMP4 ligands. *Right:* Illustration of mouse embryo with two cells expressing mis-localized LTG mutant BMP receptors, leading to ectopic pSMAD1/5 activity. A and P denote anterior and posterior, respectively. **b**, Confocal image of a hESC transfected with mutant receptor plasmid (BMPR1A^A514G^-Clover-IRES-BMPR2^A494G^), immunostained for pSMAD1/5 after a 30 min BMP4 induction. *From top:* DAPI, ZO-1, Clover (LTG mutant receptors), pSMAD1/5, and color-combined channels. Scale bar 10 μm. **c**, pSMAD1/5 intensities of hESCs after 30 min BMP4 induction: cells at edge of colony (Edge, n=23); non-edge cells expressing mutant BMPR1A^A514G^ and BMPR2^A494G^ receptors (LTG, n=73); non-transfected neighbors of transfected cells (Neighbor, n=166); cells expressing mutant receptors but without BMP4 induction (LTG no induction, n=96); and cells transfected with GFP plasmid (n=51). Z-score denotes number of standard deviations beyond background mean (of neighboring non-transfected cells). **d**, E6.25 mouse embryo transfected with mutant receptor plasmid (BMPR1A^A514G^-Clover-IRES-BMPR2^A494G^), immunostained for ZO-1, Clover (LTG mutant receptors), and pSMAD1/5. Image shows localization of mutant receptors at both apical and basolateral membrane and pSMAD1/5 activity in a transfected cell. Brown arrows indicate transfected cells. **e**, pSMAD1/5 intensity epiblast cells: cells at edge of epiblast (Edge, n=42); non-edge cells transfected with mutant receptor plasmid (LTG, n=27); their neighboring non-transfected cells (Neighbor, n= 52); and cells transfected with a GFP plasmid (GFP, n=20). Edge and LTG cells with z-score greater than 25 are not shown (n=26). Scale bar 20 μm.

We next explored whether BMP receptors are similarly localized in the basolateral membrane of mouse epiblast cells *in vivo*. To visualize receptors specifically on the cell membrane, we developed a protocol for surface-immunostaining the mouse epiblast around the start of gastrulation (see METHODS). After collection of E6.5 mouse embryos, we surgically removed the ExE from each embryo and exposed the epiblast to BMPR1A antibodies. We subsequently fixed and permeabilized the embryos and immunostained them for tight junction protein ZO-1 and epiblast marker OCT4. Light-sheet microscopy of the immunostained embryos revealed that BMPR1A receptors in epiblast cells are localized on the basolateral membrane facing the underlying VE (Fig. 2d-h and Supplementary Fig. 4k-n).

### A robust BMP signaling gradient forms from the epiblast edge

We asked whether the predicted formation of a robust BMP signaling gradient would occur in epiblast. We first measured the distribution of phosphorylated SMAD1/5 (pSMAD1/5, the downstream effectors of the BMP signaling pathway) in epithelial hESC colonies exposed to BMP4 ligands. These epithelial colonies have impermeable tight junctions and a narrow, permeable basement membrane matrix underneath mimicking an interstitial space. The tissue geometry therefore is comparable to the geometry of the epiblast in mammalian embryos ^16,25^. Akin to the simulation, we observed pSMAD1/5 gradients organized from the edges of epithelial hESC colonies exposed to spatially uniform concentrations of BMP4 (Fig. 3a,b and Supplementary Fig. 5a,b). These BMP4 signaling gradients were robust to changes in ligand concentration: colonies exposed to BMP4 concentrations across a 1000-fold range displayed stable pSMAD1/5 gradients inward from colony edges, with the depth of the gradient varying only between 2 and 10 cell widths (Fig. 3c,d and Supplementary Fig. 5e). The formation of these robust gradients was dependent on the segregation of apical and basolateral extracellular compartments by tight junctions. When tight junctions were disturbed by a brief treatment of passaging reagent ReLeSR or calcium chelator EGTA before BMP4 induction, signal response occurred throughout hESC colonies (Supplementary Fig. 5c). Further, if hESCs were exposed to BMP4 shortly after single-cell passaging, cells that had not yet formed tight junctions with adjacent cells showed significantly higher pSMAD1/5 activity than those surrounded by tight junctions (Supplementary Fig. 5d).

We observed similar BMP signaling gradients in early mouse embryos as well. In harvested mouse embryos stained for pSMAD1/5, we observed a gradient of pSMAD1/5 activity inward from the proximal edges of the epiblast at both the pre-streak (~E6.25) and early streak (~E6.75) stages of development (Fig. 3e,f). To test whether this signaling gradient is maintained even in uniformly high concentrations of BMP4, we surgically removed the ExE from E6.5 mouse embryos, exposing the remaining epiblast-VE cup. We then soaked the cup in media containing 10 ng/mL BMP4 for 30 minutes before fixing and immunostaining for pSMAD1/5 (Fig. 3g). In these BMP-soaked embryos, the pSMAD1/5 gradient reached only a few cell widths further from the proximal epiblast edge as compared to wild type embryos (Fig. 3g,h). This restriction of BMP signaling was maintained despite the fact that the BMP4 concentration was sufficiently high to induce pSMAD1/5 activity uniformly throughout the epiblast if its basolateral surface was exposed to ligands (Supplementary Fig. 5f). In summary, our results *in vitro* and *in vivo* show that robust gradients of BMP signaling activity form inward from the edges of epithelial tissues with basolateral receptor localization.

### Mis-localization of receptors leads to ectopic BMP4 signaling

Having verified the first two predictions of the model, we next tested whether the mis-localization of BMP receptors to the apical membrane results in ectopic BMP signaling. To do so, we designed a plasmid expressing epitope-tagged mutant copies of both *BMPR1A* and *BMPR2*, in which their LTA motifs were mutated into an LTG sequence (see METHODS). Unlike the wild type receptors, these mutant receptors localized at both the apical and basolateral membranes of hESCs transfected with these plasmids (Fig. 4a,b and Supplementary Fig. 6a). The transfected hESCs, in the absence of exogenous BMP4 ligands, did not show any significant BMP signaling activity (Fig. 4c and Supplementary Fig. 6b). After BMP4 exposure, however, cells expressing the mis-localizing receptors had significantly higher levels of nuclear pSMAD1/5 than their neighboring nontransfected cells (Fig. 4b,c and Supplementary Fig. 6a). The pSMAD1/5 levels of these transfected cells were comparable to that of non-transfected cells at colony edges (Fig. 4c). Thus, while basolaterally localized wild type BMP receptors in the interior of hESC colonies were insulated from apical ligands by tight junctions, cells with mis-localized BMP receptors could sense and respond to these ligands.

To test the effect of receptor mis-localization *in vivo*, we developed a method to deliver our mutant BMP receptor plasmid to anterior and distal regions of the epiblast that do not normally show BMP signaling activity, while leaving the rest of the mouse embryo unperturbed (see Methods, Supplementary Fig. 6c). Consistent with our results in hESCs, mutant BMP receptors were localized at both the apical and basolateral membranes of transfected epiblast cells *in vivo* (Fig. 4d). This mis-localization led to ectopic BMP signaling in cells in the anterior and distal regions of the epiblast, where neighboring non-transfected cells showed no signal response (Fig. 4d,e and Supplementary Fig. 6d). pSMAD1/5 levels in electroporated cells resembled that of cells at the epiblast edge (Fig. 4e). These data support our simulation results, in which BMP4 ligands can be present throughout the pre-amniotic cavity while basolateral BMP receptors in the epiblast are insulated from these signals.

### Distance from tissue edge and distance from signal source govern patterning of epithelial tissue

In summary, our results *in silico, in vitro*, and *in vivo* demonstrate how basolateral receptor localization and embryo geometry together, through an entropic buffering mechanism, result in the formation of robust BMP signaling gradients at tissue edges. Consistently, our mathematical model argues that an epithelial cell’s distance from the tissue edge (dedge) predict the cell’s signaling response better than its distance from the source of the signal (dsource, Fig. 5a,b). Here, the predictive power is quantified by the proficiency (the mutual information shared between the coordinate of a cell and its pSMAD1/5 levels, given as a percentage out of the total information entropy of pSMAD1/5 levels ^26^). While studies in multiple model organisms have shown that d_source_ is a critical determinant of patterning ^1,7,8,12,27^ our results argue that d_edge_ could also be an important developmental coordinate for the patterning of epithelial tissues.

**Fig. 5.**
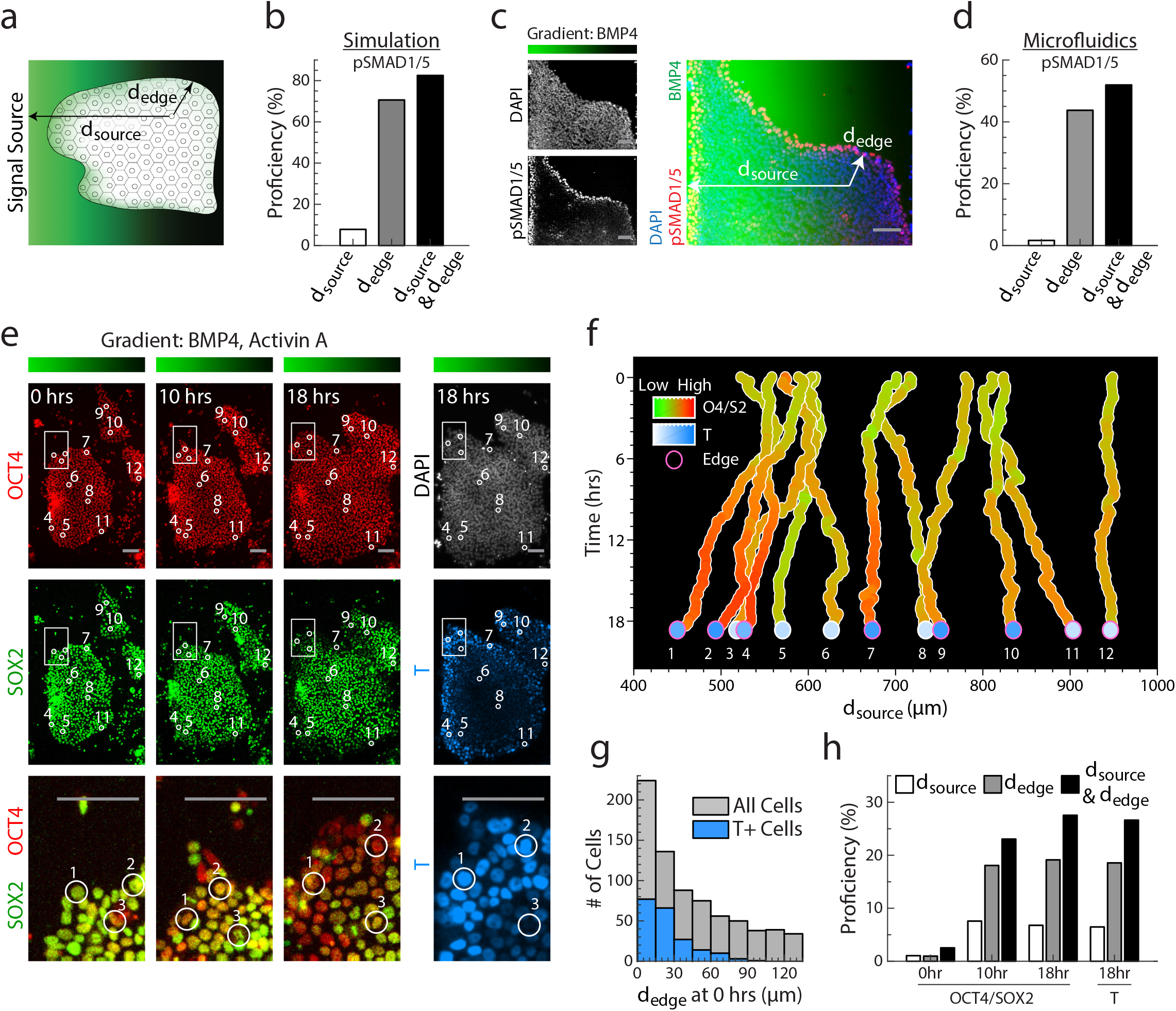
Distance from tissue edge and distance from signal source govern patterning of epithelial tissue. **a**, Illustration of epithelial tissue within a morphogen gradient emanating from a source to the left. The coordinates d_edge_ and d_source_ denote a cell’s distance from the nearest tissue edge and from the signal source, respectively. **b**, Proficiency of d_source_, d_edge_, or both coordinates to predict pSMAD1/5 level of epiblast cells in simulation (Fig. 1). **c**, Epithelial hESC colony exposed to BMP4 gradient in microfluidic device for 30 min and stained for DNA and pSMAD1/5. BMP4 gradient ranges from 10 ng/mL (green) to 0 ng/mL (black). Coordinates d_source_ and d_edge_ are depicted for a single cell as in (a). **d**, Proficiency of d_source_, d_edge_, or both coordinates to predict pSMAD1/5 levels in hESCs exposed to microfluidic BMP4 gradient (n = 13,828 cells). **e**, OCT4-RFP (red) SOX2-YFP (green) double reporter hESCs within microfluidic gradient after 0, 10, and 18 hours of differentiation. DNA, white; BRACHYURY/T, teal. Inset highlights differentiation at colony edges. Position of 12 sample cells labeled by circles. Bar above shows BMP4 and ACTIVIN A gradient within microfluidic device, ranging from 10 ng/mL (green) to 0 ng/mL (black) of each. f, d_source_ of 12 tracked cells from (E) throughout time-lapse, colored by OCT4/SOX2 ratios (red/green) and BRACHYURY/T level (teal) at end of time-lapse. Pink circles mark cells with d_edge_ less than 52 μm at 10 hours of differentiation. **g**, Distribution of d_edge_ at start of differentiation, with teal marking cells that were BRACHYURY/T+ after 18 hours of differentiation. **h**, Proficiencies of d_source_, d_edge_, or both coordinates to predict OCT4/SOX2 ratios and BRACHYURY/T levels (n=1,275 cells). Scale bar 100 μm.

To test how epithelial cell fate decisions are organized along d_source_ and d_edge_, we developed microfluidic devices capable of producing precise morphogen gradients over epithelial hESC colonies (Fig. 5c and Supplementary Fig. 7a-c). The environment within the device mimics that of a morphogen gradient produced by a signal source at the left end of the device. We exposed hESC colonies to a BMP4 gradient from 10 ng/mL to 0 ng/mL for 30 min. Consistent with our previous results (Fig. 3 and Supplementary Fig. 5a,b), signaling activity depended strongly on d_edge_ (Fig. 5c,d and Supplementary Fig. 7d). In fact, a cell’s d_edge_ had a significantly higher proficiency than d_source_ in predicting its signaling response to the BMP gradient (Fig. 5d).

To determine how d_source_ and d_edge_ correlate with cell fate decision dynamics, we built a dualcolor OCT4-RFP SOX2-YFP hESC line, in which OCT4 and SOX2 are tagged with fluorescent proteins at their endogenous loci (Supplementary Fig. 8). OCT4 and SOX2 are co-expressed in the pluripotent state (OCT4+ and SOX2+) but are differentially regulated during mesodermal differentiation (OCT4+ and SOX2-); this differential regulation is essential for the cell’s germ layer fate choice ^28^. We then cultured epithelial colonies of this hESC line in the microfluidic device, exposing them to gradients of BMP4 and NODAL-analog ACTIVIN A (from 10 ng/mL to 0 ng/mL, of each). We measured the OCT4 and SOX2 levels of individual cells in these gradients as well as their d_source_ and d_edge_ for 18 hours using time-lapse microscopy (Fig. 5e,f and Supplementary Fig. 7e). At the end of the time-lapse, we immunostained the cells *in situ* for mesodermal progenitor marker BRACHYURY/T to determine their fate choice (Fig. 5e,f and Supplementary Fig. 7f).

We found that cells with comparable d_source_ but different d_edge_ often adopted distinct cell fates (Fig. 5e,f and Supplementary Fig. 7g). In many cases, cells near colony edges had higher BRACHURY/T and lower SOX2 levels than cells in colony interiors that had a smaller d_source_ throughout the time-lapse. Furthermore, 95% of cells that expressed BRACHYURY/T at the end of time-lapse were initially located near colony edges (dedge < 66.5 μm or approximately 5.1 cell widths, Fig. 5g), where signaling is most active at the start of differentiation (Fig. 5c). After 48 hours of exposure to BMP4 and ACTIVIN A gradients, hESCs with high BRACHYURY/T and low SOX2 levels continued to be located predominantly at the colony edges, while cells in colony interiors remained undifferentiated (Supplementary Fig. 7h,i). Like pSMAD1/5, the dependence of BRACHYURY/T on d_edge_ also requires epithelial integrity. If hESCs colonies were treated with ReLeSR during first 8 hours of differentiation, BRACHURY/T appeared in the interior of the colony (Supplementary Fig. 7j).

These data argue that the organization of BMP signaling inward from epithelia tissue edges has significant implications for cell fate decisions. Indeed, we found that d_source_ and d_edge_ each carried independent information about cells’ fate choices in the microfluidic device (Fig. 5h). Furthermore, the d_edge_ of hESCs had a significantly higher proficiency of predicting their OCT4/SOX2 and BRACHURY/T levels than their d_source_, demonstrating the importance of a cell’s distance from epithelial edges as a developmental coordinate.

## Discussion

Our results reveal that the interplay between receptor localization and embryo geometry leads to formation of a robust BMP signaling gradient. Specifically, compartmentalized geometry of the early mammalian embryo requires BMP4 ligands to diffuse through a narrow interstitial space to approach basolateral receptors. This constraint limits the time and distance a ligand can travel before being captured by receptors, which are spatially restricted. As a result, a signaling gradient naturally arises, even when ligands are present uniformly in the lumen on the apical side of the epiblast. Furthermore, through a geometry-related entropic effect, BMP4 ligands accumulate in the apical lumen. Consequently, this lumen serves as a reservoir that buffers the signaling gradient against fluctuation in BMP4 concentration. Due to this entropic buffering mechanism, the channel between ExE and epiblast that connects the apical lumen containing BMP4 and basal interstitial space, acts like a stable BMP4 source. Therefore, a robust BMP signaling gradient forms spontaneously due to compartmentalized embryo geometry and basolateral receptor localization.

Developing embryo is naturally compartmentalized by epithelial tissues within. In the early mammalian embryos for example, the extracellular space is compartmentalized into the apical lumen and the basolateral interstitial space by epiblast, ^5,6,25^ Similar compartmentalization is also observed in the zebrafish migrating lateral line primordium ^29^ and the Drosophila imaginary wing disc ^30^. Indeed, TGF-β family receptors from Drosophila to mammals contain a conserved amino acid sequence motif that governs basolateral localization of receptors, implying that receptor localization is also evolutionarily conserved. Altogether, the conservation of compartmentalized geometry and receptor localization suggests that the proposed mechanism of forming robust signaling gradient is broadly applicable in many developmental contexts.

In general, the combination of compartmentalization and receptor localization allows developing tissues to sense morphogen signal in restricted environment. Such selective sensing, as demonstrated here, can dramatically modulate morphogen signaling and downstream tissue patterning. Therefore, future studies that take into account embryo geometry and receptor localization, can shed new light on how morphogen signals pattern the embryo.

## Acknowledgement

We thank Dr. Doug Richardson at Harvard Center for Biological Imaging for technical assistance. We also thank our laboratory and Dr. Xue Fei for comments on the manuscript. This work is supported by NIH Pioneer Award.

## Author contributions

Conceptualization, Z.Z. and S.R.; Methodology, Z.Z., S.Z., and S.R.; Investigation, Z.Z., S.Z., E.L., and S.R.; Writing - Original Draft Z.Z., S.Z., and S.R.; Writing – Reviewing and Editing, Z.Z., S.Z., and S.R.; Funding Acquisition, S.R., Resources, Z.Z., S.Z., E.L., J.S.G., and S.R.; Supervision, S.R.

## Competing interests

Authors declare no competing interests.

---

**Supplementary Fig. 1.**
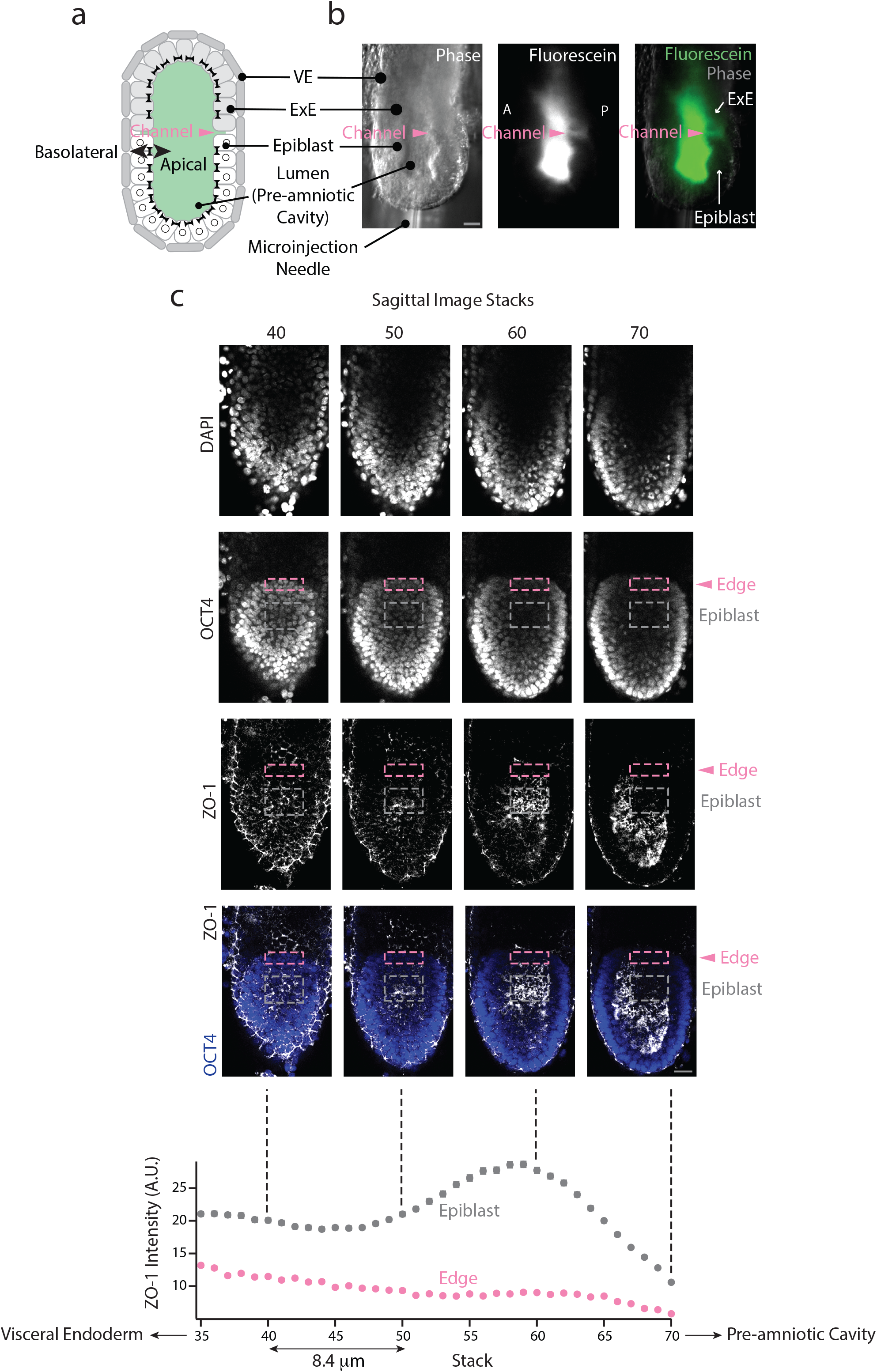
Edge of epiblast has weaker tight junctions. **a**, Illustration of pre-gastrulation mouse embryo, with the epiblast (white) and extraembryonic ectoderm (ExE, light grey) together enclosing the pre-amniotic cavity. Apical membranes of epiblast cells face the pre-amniotic cavity whereas basolateral membranes face the visceral endoderm (VE, gray). **b**, Phase, fluorescence, and color-combined images of an E6.5 mouse embryo after microinjection of fluorescein into pre-amniotic cavity. Epiblast is impermeable, but fluorescein diffuses through the gap at the edge of epiblast (border between epiblast and ExE, pink arrow). Scale bar 20 μm. **c**, *Top:* Four sagittal sections of an E6.5 embryo stained for DNA, epiblast marker OCT4, and tight junction marker ZO-1. Dotted boxes denote different areas of epiblast, in which ZO-1 intensity was quantified. *Bottom:* Average ZO-1 intensity of the edge (pink) and the rest of epiblast (gray). Scale bar 20 μm.

**Supplementary Fig. 2.**
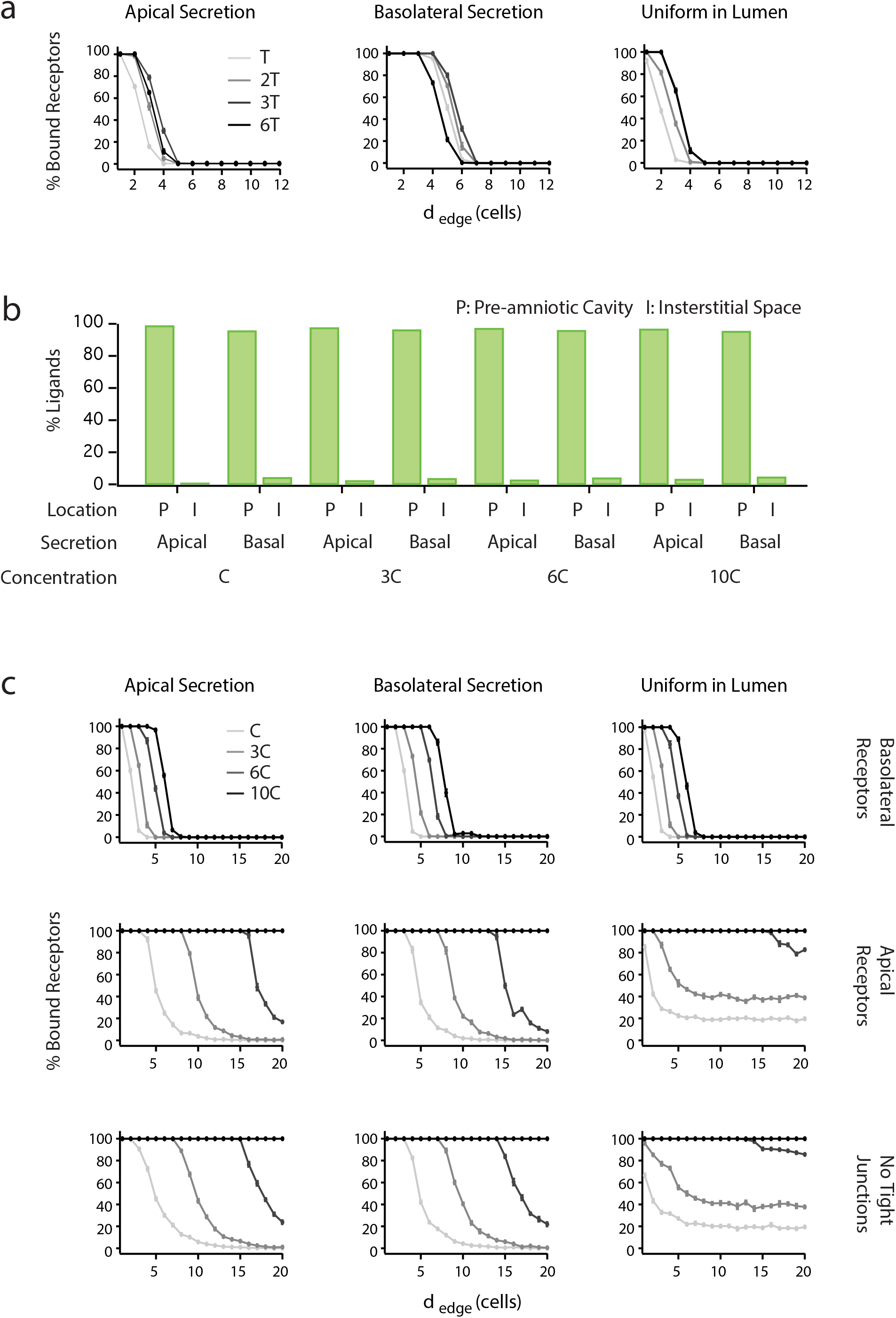
Formation of robust BMP signaling gradient is insensitive to BMP4 secretion. **a**, Percentage of ligand-bound receptors as function of distance from epiblast edge, d_edge_, over time in simulations where ligands are secreted apically (left), basolaterally (middle), or presented uniformly in lumen (right). T=5 min. **b**, Percentage of unbound ligands in pre-amniotic cavity (P) and interstitial space (I) at steady state (6T) in simulations where ligands are secreted apically or basolaterally from ExE. Here, C=0.12 ng/mL. **c**, Percentage of ligand-bound receptors as function of d_edge_ at different ligand concentration in simulations where ligands are secreted apically (left), basolaterally (middle), or presented uniformly in lumen (right). Different rows correspond to simulations where receptors are localized basolaterally (top), apically (center), or tight junctions are absent (bottom).

**Supplementary Fig. 3.**
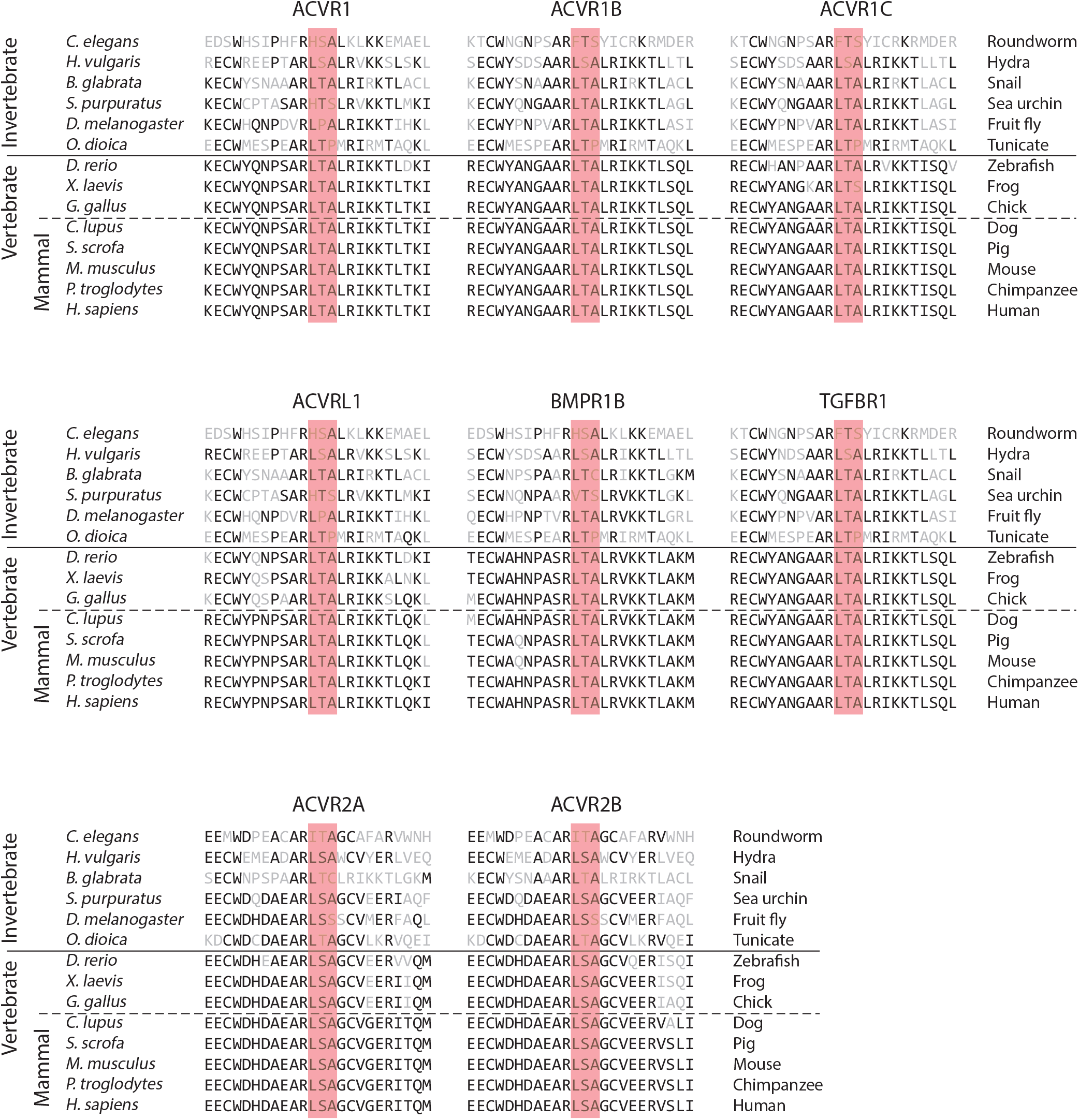
LTA motif in other TGF-β superfamily receptors. Protein sequence alignment of TGF-β superfamily receptors shows conservation of LTA motif in 6 receptors and LSA in 2 receptors.

**Supplementary Fig. 4.**
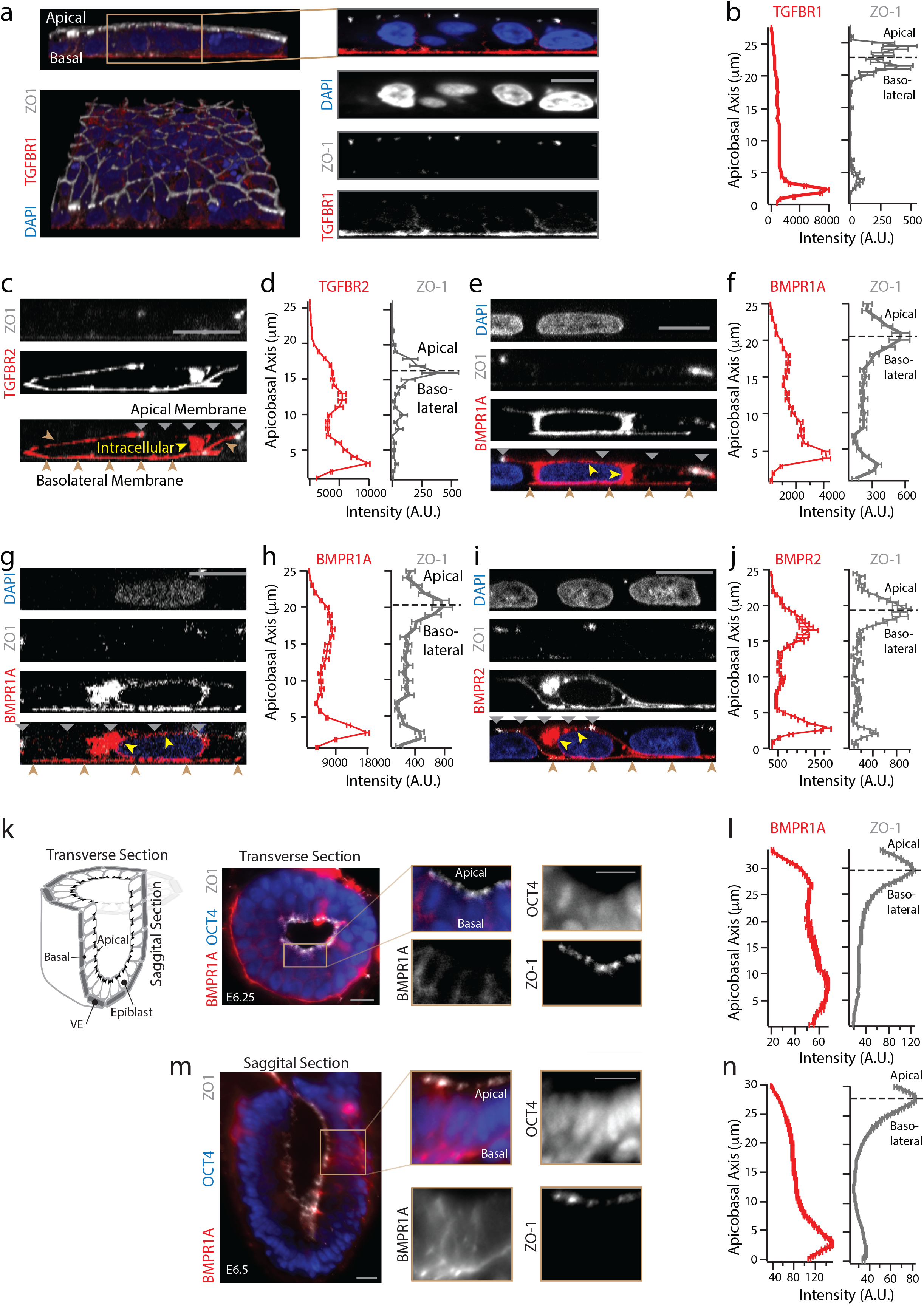
TGF-β and BMP receptors localize at basolateral membrane of hESCs in vitro and mouse epiblast in vivo. **a**, *Left Column:* 3D confocal image of hESC colony stained for DNA (blue), TGFBR1 (red), and ZO-1 (white) in lateral (top) and tilted view (bottom). *Right Column:* Zoomed-in section. Scale bar 10μm. **b**, Plots of TGFBR1 (left) and ZO-1 levels (right) against apicobasal axis show that TGFBR1 is localized below tight junctions (n=51 cells). **c**, Confocal image of a hESC expressing TGFBR2-Clover (red), stained for ZO-1 (white). Scale bar 10μm. **d**, Plots of TGFBR2-Clover (left) and ZO-1 levels (right) against apicobasal axis (n=4 cells). **e,f**, same as (**c,d**) except for BMPR1A-Clover (n=2 cells). **g,h**, same as (**c,d**) except for BMPR1A-HA (n=3 cells). **i,j**, same as (**c,d**) except for BMPR2-Clover (n=3 cells). Yellow arrows in (**c,e,g,i**) denote intracellular receptors in secretory pathway. Grey and brown arrows indicate apical and basolateral membrane. **k**, Transverse section of an E6.25 mouse embryo stained for OCT4 (blue), BMPR1A (red), and ZO-1 (white) show receptors localized at basolateral membrane of epiblast. **l**, Plots of BMPR1A (left) and ZO-1 (right) levels along apicobasal. **m,n**, same as (**k,l**), except for a sagittal section of an E6.5 mouse embryo.

**Supplementary Fig. 5.**
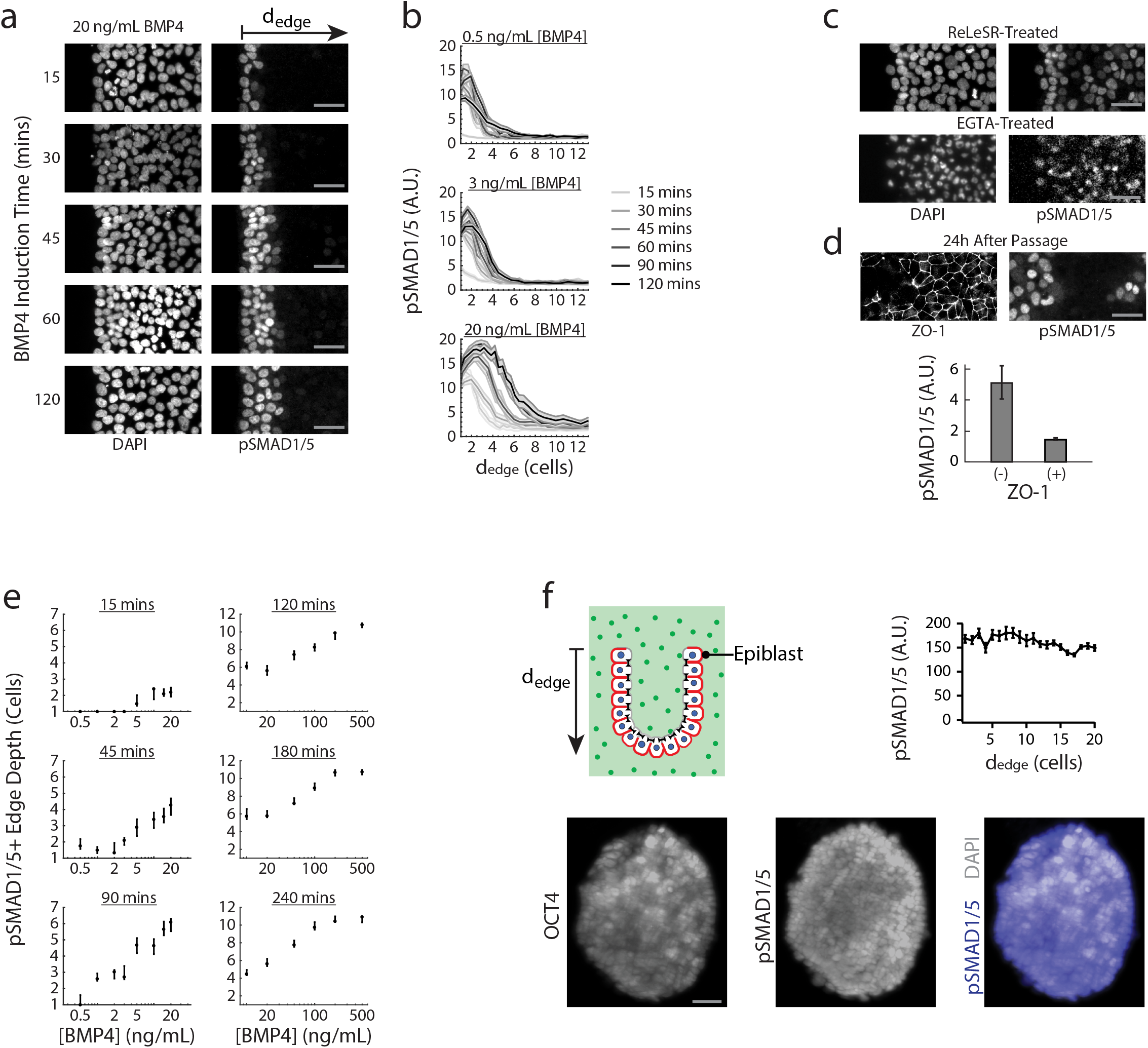
Dependence of BMP signaling gradient on epithelial integrity and embryo geometry. **a**, hESC colonies treated with 20 ng/mL BMP4 for 15-120 min and stained for DNA and pSMAD1/5. Scale bar 50 μm. **b**, pSMAD1/5 levels of hESCs as a function of their distance from the nearest colony edge, d_edge_, after 15-120 min of BMP4 treatment. **c**, hESCs stained for DNA and pSMAD1/5 after ReLeSR (top) or EGTA (bottom) treatment (see METHODS) followed by 20 min BMP4 induction. Scale bar 50 μm. **d**, *Top:* hESCs exposed to BMP4 for 20 min and stained for ZO-1 and pSMAD1/5 at 24 hours after single cell passage. *Bottom:* Average pSMAD1/5 levels of hESCs surrounded by tight junctions (+) and hESCs not surrounded by tight junctions (-), as indicated by ZO-1 immunostain. hESCs are exposed to 5 ng/mL BMP4. **e**, pSMAD1/5 levels of hESCs as a function of their distance from the nearest colony edge, d_edge_, after 15-120 min of BMP4 treatment. **f**, *Top:* Illustration of mouse epiblast after removal of ExE and VE (see METHODS), soaking in media containing 10 ng/mL BMP4. *Bottom:* E6.25 mouse embryo stained for OCT4 and pSMAD1/5 shows BMP4 concentration is sufficient to induce signaling activity in all epiblast cells if ExE and VE are removed. *Top right:* Average pSMAD1/5 level of epiblast cells as function of d_edge_ in soaked embryos with ExE and VE removed. Error bars denote SEM and scale bar 20 μm.

**Supplementary Fig. 6.**
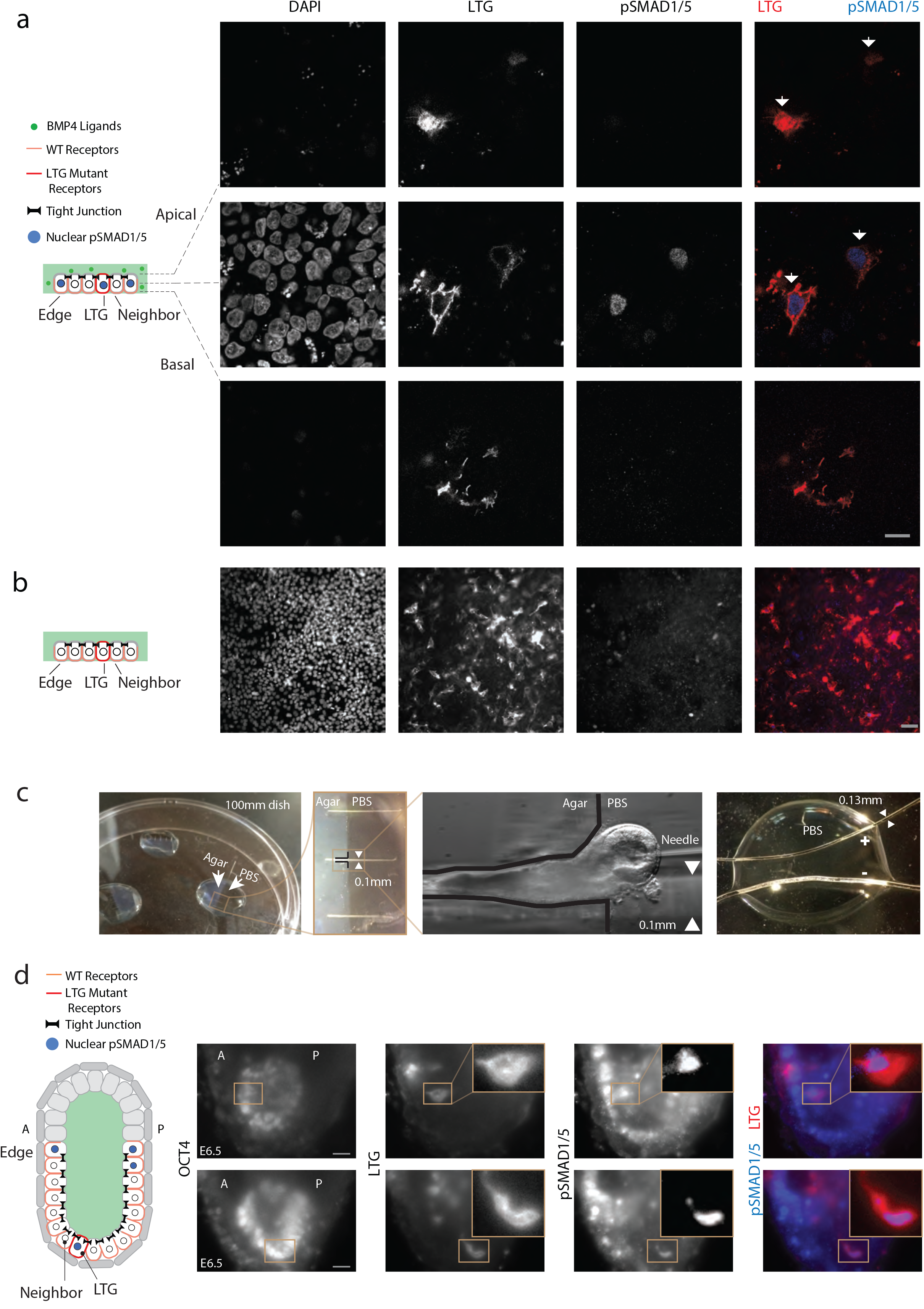
Receptor mis-localization *in vitro* and *in vivo*. **a**, Apical (top), middle (middle), and basal (bottom) image stacks of hESC colony transfected with mutant receptor plasmid (BMPR1A^A514G^-Clover-IRES-BMPR2^A494G^) and exposed to 10 ng/mL BMP4 for 30 mins. Two cells in center of colony expressed mis-localized BMP receptors. *(Left to Right):* DAPI, Clover (LTG mutant receptors), pSMAD1/5, and color-combined channels. These images show that receptor mis-localization leads to ectopic pSMAD1/5 activation. Scale bar 10μm. **b**, hESC colony transfected with mutant receptor plasmid without BMP induction. *(Left to Right):* DAPI, Clover (LTG mutant receptors), pSMAD1/5, and color-combined channels. These images show that the expression of mutant receptors in the absence of BMP4 does not result in ectopic pSMAD1/5 activation. Scale bar 40μm. **c**, Custom-made device for microinjection (left) and electroporation (right, see METHODS). **d**, *Left:* Illustration of mouse embryo transfected with mutant receptor plasmid (BMPR1A^A514G^-Clover-IRES-BMPR2^A494G^). *Right:* Two transfected E6.5 mouse embryos stained for OCT4 and pSMAD1/5 (blue), showing that cells expressing mutant receptors (red) have ectopic pSMAD1/5 activity. Scale bar 20μm.

**Supplementary Fig. 7.**
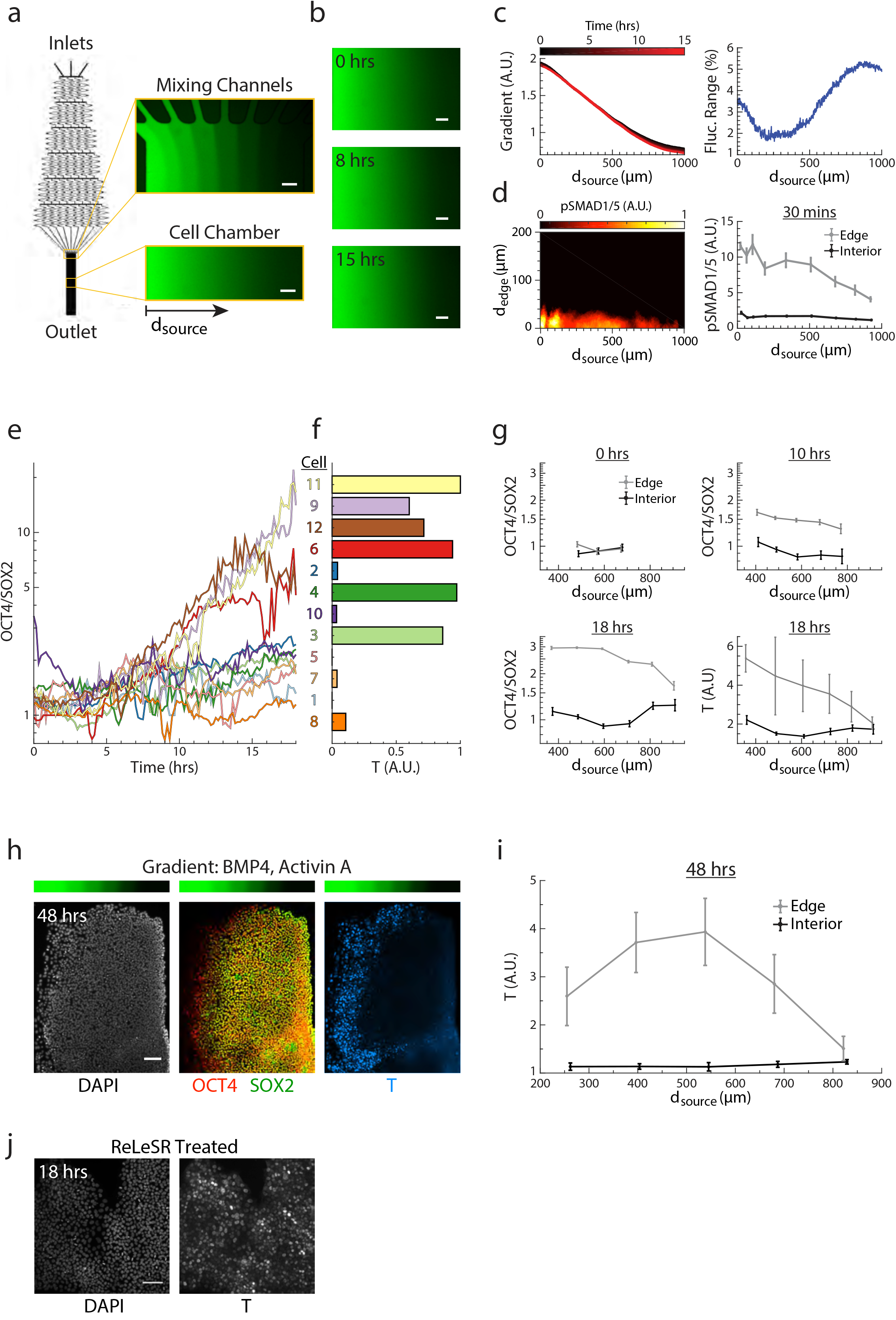
Distance from tissue edge and distance from signal source govern patterning of epithelial tissue. **a**, Diagram of microfluidic device. (Inset) Visualization of gradient using fluorescein at top and middle of cell chamber in microfluidic device. **b**, Visualization of microfluidic gradient after 0, 8, and 15 hours of flow. **c**, Level of fluorescein gradient over 15 hours (left) and its maximum range of fluctuation (right) across cell chamber as function of distance from left side of the chamber (dsource). Fluctuation range is given as percentage of total gradient range. **d**, (Left) Heat map of pSMAD1/5 levels of hESCs exposed to BMP4 gradient for 30 min as a function of d_source_ and d_edge_. (Right) pSMAD1/5 levels of edge and interior hESCs as a function of d_source_. Edge cells are defined as cells within 26 μm of a colony edge, while interior cells are those further than 78 μm from the nearest colony edge. The response of cells at the edge of the colony decreases with d_source_. **e**, OCT4/SOX2 ratios of the 12 tracked cells from Fig. 5e over time-lapse experiment. **f**, T levels of 12 tracked cells from Fig. 5e at end of time-lapse experiment. **g**, OCT4/SOX2 ratios at 0, 10, and 18 hours and T levels at 18 hours during time-lapse differentiation of edge and interior hESCs as a function of d_source_ (n=1,275 cells). **h**, Epithelial OCT4-RFP SOX2-YFP hESC colony after 48 hours of differentiation of BMP4 and ACTIVIN A gradient (above) as in Fig. 5e, stained for DNA and T. **i**, T levels of edge and interior hESCs after 48-hour gradient differentiation as a function of d_source_. Here, edge cells are defined as cells within 126 μm of colony edge, while interior cells are those further than 283 μm from nearest colony edge. Error bars denote 95% confidence intervals and scale bar 100 μm. **j**, ReLeSR treated hESCs were exposed to uniform BMP4 and ACTIVIN A (10 ng/mL of each) for 18 hours, and then stained for DNA and BRACHYURY/T. hESCs were treated with ReLeSR three times during the first 8 hours of differentiation. Scale bar 100 μm.

**Supplementary Fig. 8.**
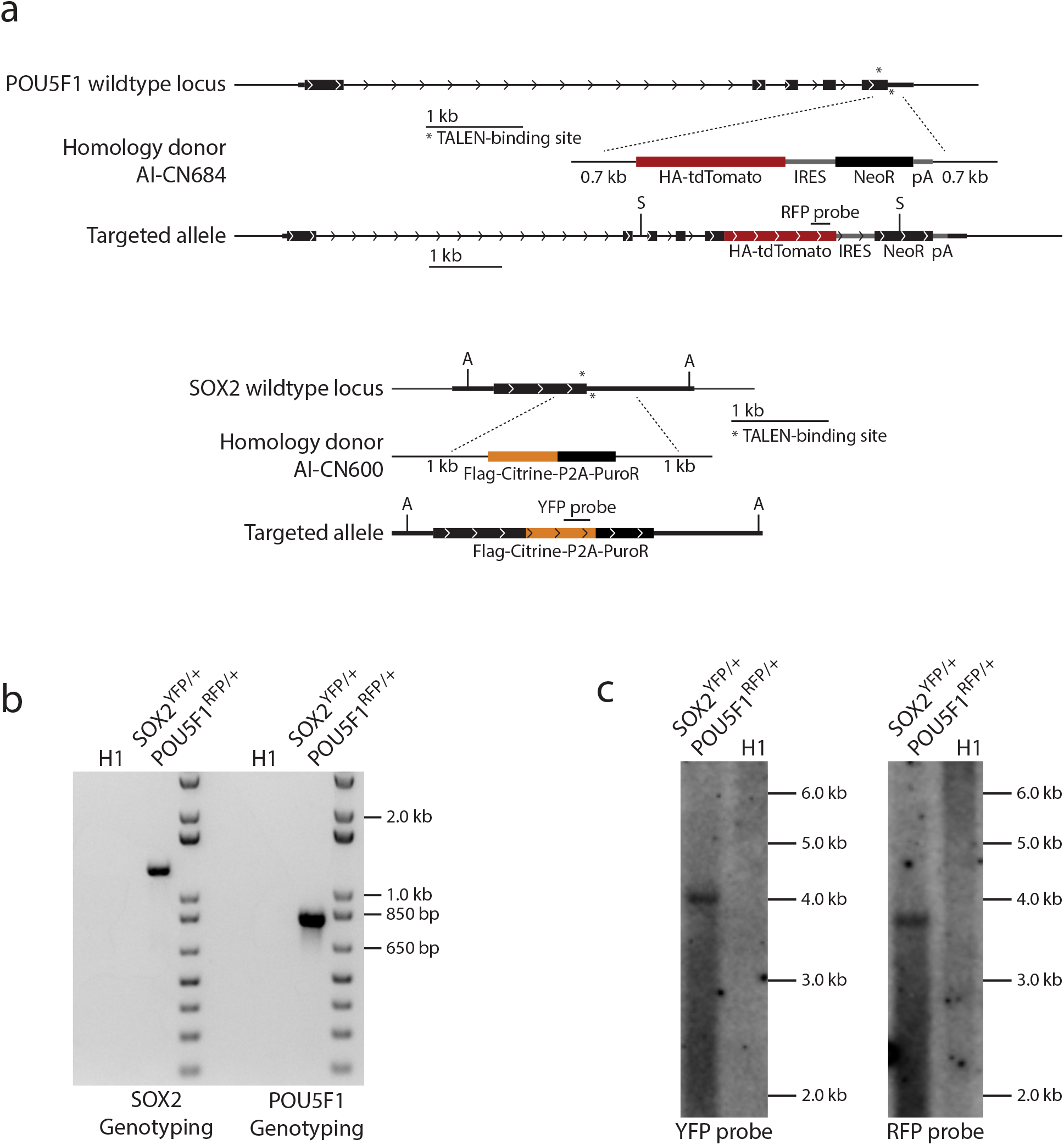
Transgenic *SOX2^YFP/+^POU5F1^RFP/+^* H1 human embryonic stem cell reporter lines. **a**, Targeting strategy to generate *SOX2^YFP/+^POU5F1^RFP/+^* reporter cell line. **b**, 5’ junction genotyping PCR of the resulting line. **c**, Southern blots of the resulting line. Restriction enzyme sites (A, AflII; S, SacI) are indicated. POU5F1 is also known as OCT4.

## Methods

### Simulation of BMP4 dynamics

BMP4 ligands were secreted by 6 ExE cells, and received by 20 epiblast cells. Each cell was 10 μm wide and 20 μm tall. The pre-amniotic cavity above the cells was 260 μm wide and 30 μm tall. The interstitial space was 260 μm wide and 2 μm tall. The lateral separation between cells was 2 μm. The simulation setup is therefore comparable to geometry of a pre-gastrulation embryo (300 μm long and 100 μm width). The height of interstitial space and lateral separation (2 μm) were estimated based on fluorescein injection experiments and embryos stained for BMPR1A. The total number of receptors was 2000, distributed uniformly between epiblast cells. Initially, 300-3000 ligands were secreted uniformly from ExE, either at the apical membrane or the basal membrane. After secretion, ligands diffusion was simulated as a random walk using Langevin dynamics ^18^. The step size along each axis was generated by random number function ran2 and displacement followed <x^2^> = 2Dt, where D = 60 μm^2^/s is the diffusion coefficient. This is based on the measurement of the diffusion of TGF-β ligand NODAL in zebrafish ^31^. The diffusing ligands were not allowed to pass tight junctions, cell membranes without receptors, or the boundaries of pre-amniotic cavity and interstitial space. Instead, these incoming ligands would be reflected at those surfaces. Given that tight junctions are absent between the ExE and the epiblast, ligands in the pre-amniotic cavity were allowed to reach the interstitial space, and vice versa through the gap at the edge of epiblast. Ligands were captured by receptors if they contacted the basolateral membrane of an epiblast cell with unbound receptors. Each epiblast cell originally had a total of 100 unbound receptors. Each lateral membrane had 40 receptors while the basal membrane had 20 receptors. Once all receptors on a membrane were ligand-bound, the membrane could no longer accept new ligands. After 15 minutes, a timescale related to endocytosis and recycling of receptors ^32,33^, each receptor-ligand pair was replaced by an unbound receptor at the same epiblast cell and an unbound ligand released by the same ExE cells. The simulation had no other parameters, and was coded in C.

### Cell lines used in the study

All human embryonic stem cell (hESC) experiments were performed with WA01 (H1) cells or SOX2-YFP, OCT4-RFP double reporter cells (see below) in a H1 background.

### Cell culture and passage

hESCs were maintained in the feeder-free cell culture medium mTeSR1 (STEMCELL Technologies) with daily media changes. For passaging, cells were dissociated *en bloc* with ReLeSR (STEMCELL Technologies) following the manufacturer’s protocol, and detached ES cell clumps were broken into smaller pieces (10–20 cells) by tapping the plate or gently pipetting several times with a wide-bore P1000 micropipette (Corning). Cells were passaged at a 1:12 split ratio onto Matrigel-coated (Corning) plates. Immediately following passage, cells were maintained in mTeSR1 supplemented with 10 μM ROCK inhibitor Y-27632 (STEMCELL Technologies) for 24 hours before returning to culture in mTeSR1 alone.

### Surface immunostaining of hESCs

Before surface receptor staining ^20^, cells were rinsed once in 1X PBS (Lonza). Cells were incubated with primary antibodies diluted in mTeSR1 with 1% BSA and 5% normal donkey serum, at 37 °C for 45 minutes. Afterward, cells were rinsed two times in PBS and subsequently fixed in 4% formaldehyde for 20 min at room temperature. See next section for secondary stains.

### Intracellular immunostaining of hESCs

Cells were fixed for 20 min at room temperature in 4% formaldehyde and rinsed three times with PBS. Permeabilization and blocking were performed simultaneously by incubating cells in blocking buffer (PBS with 5% normal donkey serum and 0.3% Triton X-100) for 60 min at room temperature. Primary antibody incubation was performed overnight at 4 °C in antibody dilution buffer (PBS plus 1% BSA, and 0.3% Triton X-100). The next day, cells were washed with PBS three times and then incubated with DAPI and secondary antibodies in antibody dilution buffer (as above) for 1 hour at room temperature. After secondary stain, cells were washed with PBS three times before imaging.

### Antibodies

BMPR1A (1:20, sc20736, Santa Cruz)

BRACHYURY/T (1:400, AF2085, R&D)

Clover (1:600, EMU101, Kerafast)

OCT4 (1:800, sc8628, Santa Cruz) pSMAD1/5 (1:800, 13820s, Cell Signaling)

TGFBR1 (1:20, sc9048, Santa Cruz)

ZO-1 (1:100, 33-9100, Thermo Fisher)

ZO-1-FITC (1:100, 33-9111, Thermo Fisher)

### Plasmid construction and transient expression of receptors

Receptor genes (BMPR1A and BMPR2) were cloned into the plasmid pCAGIP-TGFBR2-Clover (a gift from Jeff Wrana lab at Lunenfeld Tanenbaum Research Institute) between restriction sites XhoI and AgeI. To visualize receptors using small epitope tags, Clover was replaced by Myc tag or HA tag between restriction sites AgeI and NotI. To minimize side effect caused by plasmid expression of tagged protein, we excluded cells with excessive level of expression, protein aggregation of fluorescence proteins, and membrane blebbing.

### Plasmid construction and receptor mis-localization

To mis-localize receptors, LTA motifs in both *BMPR1A* and *BMPR2* were mutated into LTG sequences ^21^ by site-directed mutagenesis (NEB). The puromycin in the pCAGIP-BMPR1A-Clover plasmid was replaced by BMPR2-Myc between restriction sites BmgBI and SacI. To minimize side effect caused by plasmid expression of tagged protein, we exclude cells with excessive level of expression, signs of protein aggregation induced by fluorescence proteins, or membrane blebbing.

### hESC transfection

Transfection of hESCs was performed using jetPrime (Polyplus-transfection) or the Amaxa Nucleofector II (Lonza). For jetPrime transfection, hESCs were transfected within 2 days after passage, following the manufacturer’s protocol. For nucleofection, hESC cell colonies were dissociated into single cells (see Single cell passaging) and split into aliquots of 800,000 cells. Aliquots were spun for 3 minutes at 200 x g before resuspension in 82 μL human stem cell Nucleofector Solution 2 (Lonza) and 18 μL Supplement 1 (Lonza) with 1 - 5 ug of DNA. The cell suspension was added to a nucleofection cuvette and transfection was carried out using nucleofection program B016. Immediately following transfection, 500 uL of mTeSR1 culture medium (STEMCELL Technologies) supplemented with 10 μM ROCK inhibitor (STEMCELL Technologies) was added to the cuvette, and cells were seeded into a 15 mm well (Corning) coated with Matrigel (Corning).

### Breaking tight junctions

hESCs colonies were washed once with PBS, then treated with ReLeSR (STEMCELL Technologies) for 1-2 minutes at 37 °C. Alternatively, cells were washed once with PBS, then treated with 2mM EGTA (SIGMA) for 20 minutes ^34^ at 37 °C.

### Single cell passaging

hESCs colonies were dissociated into single cells by adding 1 mL of 0.25 % Trypsin-EDTA (Life Technologies) or 1 mL Accutase (Innovative Cell Technologies) to cells in a 9.6 cm^2^ well, incubating cells for 5-7 mins at 37 °C, and quenching with 1 mL of ES-qualified FBS (Millipore). Cell clumps were broken up by gently flushing cells 5-10 times with a P1000 micropipette. Afterward, cells were collected, centrifuged at 200 x g for 3 m, and re-suspended in mTeSR1 supplemented with 10 μM ROCK inhibitor. 200,000 to 1,200,000 cells were seeded into a 15 mm well coated with Matrigel.

### Epifluorescence imaging of hESCs

hESCs were imaged on a Zeiss Axiovision inverted microscope with Zeiss 10× and 20× plan apo objectives (NA 1.3) using the appropriate filter sets and an Orca-Flash 4.0 camera (Hamamatsu). The 38 HE GFP/43 HE DsRed/46 HE YFP/47 HE CFP/49 DAPI/50 Cy5 filter sets from Zeiss were used.

### Confocal imaging of hESCs

Cells were imaged on a Zeiss LSM 700 confocal microscope with Zeiss 40× and 63× oil objectives (NA 1.3) with the appropriate filter sets and a back-thinned Hamamatsu EMCCD camera.

### Mouse embryo recovery

Eight weeks old adult C57BL/6J female mice were naturally mated and sacrificed at 6am (E6.25), 12pm (E6.5), or 6pm (E6.75) on the 6th day post coitum. In each case, the uterus was recovered and embryos were dissected from the deciduae ^35,36^ in embryo culture buffer (see below).

### Mouse embryo microinjection

Embryos were transferred to a microinjection chamber immersed in PBS. These microinjection chambers were made with 0.4% agarose and had multiple channels for holding embryos (Supplementary Fig. 6c). They were specifically designed to minimize the movement and deformation of embryos during microinjection. Microinjection needles were made by pulling glass capillaries (Kwik-Fil, 1B100F-4, World precision instruments) in a micropipette puller (Model P-97, Sutter instrument) using a custom program (Heat 516, Pull 99, Vel 33, and Time 225). The needle was back-filled with 1.5-2.0 μg/μL plasmid purified using endotoxin-free maxiprep kit (NucleoBond Xtra Maxi Plus EF, 740426.10, Macherey-Nagel). To reduce jamming during microinjection, the plasmid solution was centrifuged at 5,000g for 10 min, and the supernatant was loaded into the needle. The microinjection needle was inserted into the pre-amniotic cavity and the plasmid solution was injected using air pressure (XenoWorks digital microinjector, Sutter instrument) so that the cavity expanded slightly.

### Mouse embryo electroporation

Microinjected embryos were transferred to electroporation chamber immersed in PBS (Supplementary Fig. 6c). Electrodes in the chamber were made of 0.127 mm platinum wires (00263, Alfa Aesar). Embryos were placed at the center of the chamber, either parallel or perpendicular to platinum wires. Three electric pulses ^37^ (30 V, 1 ms duration, 1 s apart) were delivered using electro square porator (ECM 830, BTX).

### Mouse embryo culture

Electroporated embryos were transferred to 12-well cell culture dish containing embryo culture media at 37 °C and 5% CO2. This media ^38^ contains 50% rat serum (AS3061, Valley Biomedical) and 50% Ham’s F12 (31765035, Thermo Fisher) supplemented with N-2 (17502048, Thermo Fisher). The media was equilibrated in the incubator for 1 hour prior to embryo addition. E7.5 embryos cultured in this media developed heartbeats after 24-36 hours (Video S1). Electroporated E6.5 embryos were cultured for 4 hours. Only embryos without visible defects were subjected to downstream analysis.

### Surface immunostaining of embryos

Extraembryonic ectoderm and underlying visceral endoderm were removed using fine forceps (1125200, Dumont). The remaining epiblast and visceral endoderm were incubated in primary antibodies diluted in embryo culture media with 1% BSA and 5% normal donkey serum for 45 min at 37 °C and 5% CO2. The embryos were subsequently washed three times with PBS, before fixed for 30 min at room temperature with 4% formaldehyde. Due to this fixation step, occasionally aggregates of unbound antibodies were retained inside the pre-amniotic cavity. These large aggregates with no DAPI or OCT4 stain, were excluded from analysis.

### Intracellular immunostaining of embryos

Embryos were fixed for 30 min at room temperature in 4% formaldehyde and rinsed 3 times with PBS. Permeabilization and blocking were performed simultaneously by incubating cells in 5% normal donkey serum, 1% BSA, and 0.3% Triton X-100 in PBS for 1 hour at room temperature. Primary antibody incubation was performed overnight with 1% BSA, and 0.3% Triton X-100 in PBS at room temperature. In the morning following primary incubation, embryos were washed 3 times with PBS then incubated with secondary antibodies in staining buffer (as above) for 1 hour at room temperature. After secondary stain, embryos were washed three times with PBS before imaging.

### Light-sheet imaging of embryos

Stained embryos were embedded into low-melting agarose (BP165-25, Thermo Fisher) containing 0.1 μm fluorescent beads (F8801, Thermo Fisher). The embedded embryos were then imaged in Zeiss Light-sheet Z1 microscope under 20x water objective from 4 angles. The resulting multi-view images were registered using ImageJ plugin multi-view reconstruction.

### Fabrication of microfluidic devices

Microfluidic devices were fabricated in poly(dimethylsiloxane) PDMS using rapid prototyping and soft lithography following published procedures ^39^. A photomask was designed based on published work to create microfluidic devices that generate linear concentration gradients. A 100 μm thick “negative” master mold was fabricated from the photomask by patterning SU-8 3050 photoresist on an Si wafer through photolithography. “Positive” replicas were generated by molding PDMS against the master. After devices were cured, three inlets and one outlet with 0.5 mm diameters were punched. The mold-side surfaces of devices were rendered hydrophilic by plasma oxidation through a 5-minute plasma treatment in room air with a plasma cleaner (Harrick Plasma) at high RF power. Immediately after plasma treatment, devices were submerged in deionized water and autoclaved at 121 degrees Celsius and 100 kPa for 20 minutes in liquid cycle to simultaneously sterilize the devices and remove toxic non-cross-linked monomers. Bubbles were removed from the autoclaved devices by vacuum desiccation for 30 minutes. Afterward, autoclaved Tygon tubing (Saint Gobain) was inserted into inlets and outlets, and the entire device was sterilized again with 30 minutes of UV light in a Class II Biological Safety Cabinet. For all experiments using the microfluidic devices, the amount of time the microfluidic devices spent not submerged in water or cell culture media after plasma treatment was minimized to maintain the hydrophilicity of the molded surface.

### Culture of hESCs in microfluidic devices

hESCs to be cultured in microfluidic devices were passaged and maintained in dish culture as described earlier in METHODS. At 1 hour prior to application of microfluidic devices, cell culture media was changed to mTeSR supplemented with penicillin-streptomycin solution (100X, Sigma-Aldrich). Immediately prior to application of microfluidic devices, the tubing of microfluidic devices was filled with mTeSR + penicillin-streptomycin and clamped closed at ends. Devices were then directly attached to the hESC dish using an aluminum clamp designed to fit the dish. Microfluidic devices were positioned with their molded surface over the hESCs and gently clamped downward onto the dish such that cells were located in the cell chamber. Afterward, inlet tubing was connected to media reservoirs containing mTeSR + penicillin-streptomycin, and outlet tubing was connected to a 3 mL syringe loaded on a syringe pump (Harvard Apparatus). The syringe pump was set to withdraw fluid at a flow rate of 20 μl/min or less. The clamped dish was then placed back into an incubator or loaded onto a Zeiss Axiovision inverted microscope for time-lapse imaging, followed by unclamping all attached tubing and starting the syringe pump. After an hour of flow through the microfluidic device to prime the gradient over the cells, the media in reservoirs was changed to the appropriate differentiation conditions either by adding chemicals directly or by progressive dilution. At the end of microfluidic experiments, 1 mg/mL fluorescein isothiocyanate-dextran (Sigma Aldrich) was added to inlet reservoirs to measure the gradient profile within the device. Once a stable gradient was detected and imaged, the microfluidic device was unclamped from the plate, and cells were fixed and immunostained *in situ* following procedures described in intracellular staining of hESC.

### Construction of dual-color hESCs

TALEN genes targeting *POU5F1* (AI-CN330 targeting TCTGGGCTCTCCCAT; AI-CN331 targeting TCCCCCATTCCTAGAAGG) were prepared using the REAL method (PMID: 21822241) to match reported target sites (PMID: 21738127). The TALEN genes targeting *SOX2* (AI-CN298 targeting TTAACGGCACACTGCCC; AI-CN299 targeting TCCAGTTCGCTGTCCGGC) were made by the Joung lab (Massachusetts General Hospital) using the FLASH method (PMID: 22484455). *POU5F1* homology-directed repair (HDR) donors AI-CN623 and AI-CN684 were used for constructing the *POU5F1^RFP/+^* and *SOX2^YFP/+^POU5F1^RFP/+^* lines, respectively. The *SOX2* HDR donor was AI-CN600.

H1 hESCs (WA01; WiCell) were maintained with mTeSR1 media (Stem Cell Technologies) on Matrigel (Corning). Stem cells at p38-39 were treated with 1 μM thiazovivin (StemRD) one day prior to electroporation (Neon; Invitrogen; resuspension buffer R; 100 μL electroporation tip; 1050 V, 30 ms pulse width, 2 pulses; 1.5 or 2 × 10^6^ cells) as single cells (StemPro Accutase, Life Technologies) with 1.5 or 3 μg of each TALEN plasmid and 6 or 12 μg of the HDR donor plasmid. The cells were treated with 2 μM thiazovivin for 24 h following electroporation. After recovery, cells were treated with 1 μg/mL puromycin (Life Technologies) for three days. Following three days of recovery, dual *SOX2* and *POU5F1*-targeted cells were treated with 75 μg/mL G418 sulfate (Life Technologies) for three days. Fluorescent colonies were validated by PCR *(SOX2* 5’ junction primers: CCTGATTCCAGTTTGCCTCTCTCTTTTTTTC, CTTATCGTCGTCATCCTTGTAATCAGATCTCC; *POU5F1* 5’ junction primers: AT GCTGTTACTCAGCAAGTCCAAAGCTTG, GCGTAGTCTGGGACGTCGTATGGGTAAG), had normal karyotypes (Cell Line Genetics), and Southern blots (Lofstrand) confirmed insertion of fluorescent protein transgenes at only the targeted loci in *SOX2^YFP/+^POU5F1^RFP/+^* (AI01e-SOX2OCT4) and *POU5F1^RFP/+^* (AI05e-OCT4RFP). Silencing of SOX2-YFP was occasionally observed in a small fraction of *SOX2^YFP/+^POU5F1^RFP/+^* cells. This silenced population was regularly removed by fluorescence-activated cell sorting (FACS).

### Time-lapse microscopy

For live-cell microscopy, a Zeiss Axiovision microscope was enclosed with an environmental chamber in which CO2 and temperature were regulated at 5% and 37 °C, respectively. Time-lapse images were acquired every 10 min for 18-48 hrs. Image acquisition was controlled by Zen (Zeiss); all cell tracking was manually performed using the TrackMate package in ImageJ (NIH). Cell segmentation and fluorescence measurements were done using CellProfiler ^40^. All other image data analysis was performed using custom code written in Matlab (MathWorks).

